# Aberrant accumulation of NIK promotes tumorigenicity by dysregulating post-translational modifications in breast cancer

**DOI:** 10.1101/2021.08.27.457878

**Authors:** Yusuke Hayashi, Jun Nakayama, Mizuki Yamamoto, Masashi Maekawa, Shinya Watanabe, Shigeki Higashiyama, Jun-ichiro Inoue, Yusuke Yamamoto, Kentaro Semba

**Author notes:** Corresponding author Jun Nakayama and Kentaro Semba, Department of Life Science and Medical Bioscience, School of Advanced Science and Engineering, Waseda University, TWIns, 2-2 Wakamatsu-cho, Shinjuku-ku, Tokyo, 162-8480, Japan.

## Abstract

Post-translational modifications and mRNA translation are frequently altered in human cancers. However, investigations to understand their roles in the cancer progression mechanism remain insufficient. In this research, we explored protein levels altered by translational or post-translational regulation by analyzing transcriptome and western blotting data of the highly malignant breast cancer cell lines. From these analyses, NIK was found to be upregulated at the protein level to predominantly activate the non-canonical NF-κB pathway in a breast cancer cell line. Furthermore, the increase in NIK protein production was attributed to the dysregulation of ubiquitin-proteasome system caused by a decrease in the translation of cIAP1. NIK upregulation contributed to tumorigenicity by regulating the expression of inflammatory response-related genes. Collectively, our study suggests that NIK is post-translationally modified and has the potential to be a therapeutic target and diagnostic marker for breast cancer.

## Introduction

Breast cancer is the most common cancer among women worldwide ^1^. Although breast cancer mortality has decreased for three decades since the 1990s, the decreasing trend in mortality has slowed in recent years ^1,2^. Breast cancer has been traditionally categorized into four molecular subtypes (luminal A, luminal B, human epidermal growth factor receptor type 2 (HER2) -enriched, and triple-negative) based on the expression of pathological marker proteins such as HER2, estrogen receptor (ER), and progesterone receptor (PR). In particular, triple-negative breast cancer (TNBC) constitutes approximately 10%∼20% of breast cancers and is characterized by defects in HER2, ER, and PR expression ^3,4^. TNBC presents a relatively poor prognosis, with metastases occurring more frequently than in other subtypes despite the limited molecular therapeutic targets ^3,5^. Therefore, it is necessary to explore innovative therapeutic targets by further elucidating the malignant mechanism of TNBC.

In normal cells, signal transductions are tightly regulated through translational and post-translational modification mechanisms, while, in cancer cells, abnormalities in translational and post-translational modifications contribute to their malignancy ^6,7^. During the translation process, oncogenic signaling activation, such as through the mTOR and RAS-MAPK pathways, induces an enhancement in eIF4F complex expression and phosphorylation, thereby promoting global translation and contributing to cancer malignancy ^6^. On the other hand, unusual tRNA modification and RNA conformations also contribute to cancer progression by promoting the translation of specific genes ^8–10^. Post-translational modifications involve a large variety of mechanisms. For example, ubiquitination, one of the major post-translational modifications, has a large number of regulatory functions, including proteostasis, signaling complex assembly, chromatin remodeling, and protein secretion ^7^. In particular, protein degradation by ubiquitination is essential for the regulation of signaling pathways, such as MAPK, NF-κB and PI3K-AKT-mTOR, which play important roles in cell growth and survival ^7^. Disruption of ubiquitin-modifying machinery can lead to the malignant transformation of cancer and a variety of other diseases ^7,11^. To elucidate mechanisms of cancer malignancy, transcriptome analysis is frequently used, and the development of analysis at single cell level has made it possible to conduct more detailed analysis ^12^. However, it is also necessary to conduct analyses integrating transcriptome analysis, translational and post-translational modification evaluations at the protein level.

*In vivo* investigations with cancer cell line-derived models have contributed to the understanding of cancer biology and cancer hallmarks ^13^. Animal models enable to analyze the functions of certain genes involved in cancer malignancy and the evaluation of the antitumor properties of preclinical candidate drugs. Among these models, xenograft models have been frequently utilized to assess tumorigenicity at particular organs and metastases to distant organs. Xenograft models are classified into two categories: orthotopic and ectopic. In particular, an orthotopic xenograft (OX) model, which mimics early cancer progression, is appropriate to comprehensively understand tumorigenesis and metastatic mechanisms. Recently we established a cell line, LM05, with high tumorigenic and lung-metastatic properties from a TNBC cell line MDA-MB-231 using an OX model ^14^. Interestingly, this cell line showed an different expression profile from that of another lung-metastatic cell line, LM1-2-1, established by tail vein injection (TVI) that corresponds to an ectopic xenograft model ^14,15^. However, the molecular mechanisms of high tumorigenic and lung-metastatic properties of LM05 cells have not yet been studied in detail.

In this research, we comparatively analyzed western blot and microarray data of LM05 cells to discover their specific activating signals in comparison with parental MDA-MB-231 cells and LM1-2-1 cells. As a result, we identified that the non-canonical NF-κB pathway was constitutively activated in LM05 cells via NIK upregulation at the protein level. The aberrant accumulation of NIK due to dysregulation of post-translational modifications promoted tumorigenicity via a cancer-inducing inflammatory response.

## Materials and Methods

### Cell culture

MDA-MB-231-*mSlc7a1*-*luc2* (parental cells), LM05 and LM1-2-1 cells were established as previously described ^14^. These cell lines were cultured in RPMI-1640 (Fujifilm Wako Pure Chemical Corporation, Osaka, Japan) supplemented with 10% heat-inactivated FBS (Nichirei Biosciences Inc., Tokyo, Japan), 100 U/mL penicillin (Meiji Seika Pharma Co., Ltd., Tokyo, Japan) and 100 μg/mL streptomycin (Meiji-Seika Pharma) at 37°C in 5% CO_2_. Plat-E packaging cells were kindly provided from T. Kitamura (Institute of Medical Science, University of Tokyo), and TIG-3 cells were kindly provided from B. Shiotani. These cells were cultured in DMEM (Fujifilm Wako Pure Chemical Corporation) supplemented with 10% heat-inactivated FBS, 100 U/mL penicillin, and 100 μg/mL streptomycin.

### Western blotting

Western blotting was performed as previously described ^16^. Cells were lysed in 1× SDS sample buffer (50 mM Tris-HCl (pH 6.8), 2% SDS, 5% 2-mercaptoethanol, 0.1% BPB, 10% glycerol) and then boiled at 95°C for 5min. To examine NIK protein level, cells were treated with 10 μM MG132 (Peptide Institute, Inc., Osaka, Japan) for 4 hours. To assess cIAP1 protein stability, cells were treated with 10 µg/mL cycloheximide (CHX) (Fujifilm Wako Pure Chemical Corporation). In addition, hypotonic buffer (10 mM HEPES-KOH (pH 7.9), 1.5 mM MgCl_2_, 10 mM KCl, protease inhibitor cocktail, and 0.15 mM DTT) was used to fractionate nuclear lysates and cytoplasmic lysates. Then, 3× SDS sample buffer was added to each lysate. The lysates were subjected to SDS-PAGE and transferred to PVDF membranes (Millipore, Darmstadt, Germany). Finally, the target proteins were detected using ImmobilonTM Western (Millipore). Information regarding the primary antibodies and secondary antibodies is described in Supplementary Table S1.

### Immunoprecipitation

Breast cancer cells were lysed in TNE buffer (20 mM Tris-HCl (pH 8.0), 150 mM NaCl, 1% NP-40, 2 mM EDTA, 25 mM NaF, 17.5 mM β-glycerophosphate, 1 mM Na_3_VO_4_, cOmplete (Roche, Basel, Switzerland)). The protein concentrations of the cell lysates were measured by the BCA method (Thermo Fisher Scientific, Waltham, MA, USA). Then, each protein lysate was added to a NIK antibody (#4994, Cell Signaling Technology, Danvers, MA, USA) or a rabbit IgG antibody (12-370, Millipore) and mixed with rotation overnight. After mixing, 20 µL of Protein A Sepharose 4 FF (GE Healthcare Japan, Tokyo, Japan) was added and mixed with rotation at 4°C for 3 hours. Subsequently, the supernatant was removed after centrifugation, and the Protein A Sepharose was washed five times with TNE Buffer. Finally, 1× SDS sample buffer was added after removing the supernatant followed by heating at 95°C for 5 minutes. This extract solution was used as the immunoprecipitation sample.

### Animal experiments

Animal experiments were performed in compliance with the guidelines of the Institute for Laboratory Animal Research, National Cancer Center Research Institute (experimental number: T18-009) and the Animal Committee of Waseda University (accession numbers: WD19-058, 2019-A068, WD20-005, 2020-A067, WD21-082, 2021-A074). Female NOD.CB-17-*Prkdc*-scid/J mice (NOD-SCID, 5–6 weeks old, Charles River Laboratories Japan, Inc., Kanagawa, Japan) were used for the orthotopic xenograft and tail vein injection models. The methods for establishing the xenograft metastasis model and tail vein injection model and performance of bioluminescence imaging were previously described ^14^. For orthotopic xenografts, a total of 1.0×10^6^ cancer cells in 10 µl of D-PBS(-) (Fujifilm Wako Pure Chemical Corporation) were implanted into the fourth fat pad of each mouse using a 28-gauge syringe. When the primary tumor volumes reached 300 mm^3^, they were resected under anesthesia with 2.5% isoflurane (Fujifilm Wako Pure Chemical Corporation). After the primary tumors were resected, mice were periodically monitored for metastasis formation using an *in vivo* imaging system (IVIS) -Lumina XRMS (Perkin-Elmer, Waltham, MA, USA) for 2 weeks. For the tail vein injection model, a total of 5.0×10^5^ cancer cells in 100 *µ*l of D-PBS(−) (Wako Pure Chemical Industries) were injected into the tail vein of each mouse using a 27-gauge needle. To monitor these xenograft mice, 200 µL of D-luciferin (15 mg/mL) (Gold Biotechnology, Inc., St. Louis, MO, USA) was injected intraperitoneally into each mouse, and bioluminescence imaging was performed with the IVIS every week. In addition, the bioluminescence of the lung metastatic tissue was measured ex vivo using the IVIS after intraperitoneal administration of 200 µL of D-luciferin to assess the metastatic potential of the cancer cells.

### Histochemical analyses

The dissected primary tumors and lung metastasis tissues from the orthotopic mouse model were fixed with 4% paraformaldehyde–PBS and embedded in paraffin. The paraffin-embedded human breast cancer cells with a breast tissue microarray (BC081116d, US Biomax, MD, USA) were baked for 2 hours at 60°C before proceeding with the following steps. The paraffin sections were deparaffinized and rehydrated in xylene, a graded ethanol series that decreased stepwise from 100% to 50%, and distilled water.

For immunohistochemistry (IHC) analysis, antigen retrieval was performed in 10 mM citrate buffer (pH 6.0) at 120°C or 95°C for 20 minutes. After cooling to room temperature, the endogenous peroxidase activity was blocked with 3% H_2_O_2_ in 10% methanol for 30 min. Then, these sections were incubated in 2.5% normal horse serum (Vector Laboratories, CA, USA) for 1hour at room temperature. The specimens were incubated with primary antibodies against α-SMA (A2547, Sigma-Aldrich Co., MO, USA), Ki-67 (ab16667, Abcam, Cambridge, UK), CD206 (AF2535, R&D Systems, Inc., MN, USA), CAM5.2 (#349205, BD Bioscience, CA, USA), and NIK (HPA027269, Sigma-Aldrich Co.) at 4°C overnight. Then, the sections were incubated with the secondary antibody solutions for 1hour at room temperature. The sections were stained with an ImmPACT® DAB EqV substrate kit (Vector Laboratories) followed by staining with hematoxylin, dehydration and mounting. Images were acquired with a BZ-X700 microscope (Keyence Corporation, Osaka, Japan) and analyzed using the image analysis application for BZ-X700 (Keyence Corporation) and ImageJ software (National Institutes of Health).

For hematoxylin and eosin (HE) staining, deparaffinized and rehydrated sections were stained in Mayer’s hematoxylin solution (Muto Pure Chemicals Co., Tokyo, Japan) for 10min. Then, the sections were soaked in 0.1% saturated lithium carbonate at 37°C for 5min. After washing in distilled H2O, the sections were stained with eosin solution (Muto Pure Chemicals Co.) for 10min. Then, the sections were immediately washed with 100% ethanol, 90% ethanol and xylene. Finally, the sections were mounted, and images were acquired with a BZ-X700 microscope.

For terminal deoxynucleotidyl transferase dUTP nick end labeling (TUNEL) staining, deparaffinized and rehydrated sections were stained with a TUNEL Assay Kit (ab206386, Abcam) according to the manufacturer’s protocols. After staining with hematoxylin, dehydration and mounting, images were acquired with a BZ-X700 microscope (Keyence Corporation).

### RNA-seq analysis

Total RNA was extracted with QIAzol (Qiagen, CA, USA) according to the manufacturer’s protocol. After purification of the total RNA, the quantity and quality of the RNA were evaluated with a Nanodrop ND-1000 spectrophotometer (Thermo Fisher Scientific). cDNA libraries for RNA sequencing were established from total RNA using NEBNext Poly(A) mRNA Magnetic Isolation Module (New England Biolabs, MA, USA) to select poly-A mRNA followed by strand-specific library preparation using MGIEasy RNA Directional Library Prep Set V2.0 (MGITech Co. Shenzhen, China). Paired-end sequencing with a lead length of 150 bases was performed on a DNBSEQ-G400 (MGI tech) platform following the manufacturer’s instructions.

The raw sequence data (fastq files) were trimmed to remove adapter sequences by Trim-Galore (v0.6.4), and then, the trimmed read data were mapped on GRCh38/hg38 (GENCODE) using HISAT2 (v2.2.0). After the SAM files were converted Bam files using SAMtool (v1.10), the gene expression FPKM was quantified by StringTie (v2.1.2). Then, read count matrices were generated for each gene by prepDE.py. After removing low-expression genes with less than 10 counts per million (CPM), differentially expressed genes were identified using edgeR in R. Statistical cutoffs based on a p-value <0.05 and a log fold change (Log2(FC)) ±1 were used to filter DEGs between LM05-shGFP cells and LM05-shNIK cells. The z scores of these DEGs were calculated by gene filtering, and hierarchical clustering heatmaps were created using pheatmap in R (v3.6.2). For pathway analysis, the expression data were analyzed with gene set enrichment analysis (GSEA; https://www.gsea-msigdb.org/gsea/index.jsp) and ingenuity pathway analysis (IPA; https://digitalinsights.qiagen.com/products-overview/discovery-insights-portfolio/analysis-and-visualization/qiagen-ipa/).

### Statistical analysis

All statistical analyses were performed with GraphPad Prism 7 (GraphPad Software, San Diego, CA, USA). Statistical results are presented as the mean ±SEM. Welch’s t test and one-way or two-way ANOVA were employed for comparisons between two or three groups of data.

### Data availability

RNA-seq data were submitted to the NCBI GEO database under accession number GSE182261. The microarray expression data of the breast cancer cells that can generate lung metastases were obtained from our previous research ^14^. These expression data and other supporting data of this research are available from the corresponding authors upon reasonable request.

Other experimental methods are included in Supplementary Methods.

## Results

### The non-canonical NF-κB pathway is constitutively activated in LM05 cells

We previously established two lung metastatic breast cancer cell lines (LM05 and LM1-2-1) ^14^. LM05 cells were established from MDA-MB-231 cells (parental cells) by two cycles of the generation of OX and subsequent extraction from the lung metastatic tissue (Fig. 1A). On the other hand, LM1-2-1 cells were established from parental cells by two cycles of TVI and subsequent extraction from the lung metastatic tissue (Fig. 1A). LM05 cells have a high potential for tumorigenicity in the fat pads compared with the parental cells and LM1-2-1 cells (Fig. 1B) ^14^. In previous studies, transcriptome analysis of these highly malignant cell lines was performed as a conventional research strategy ^17–19^. In this research, considering the importance of translational and post-translational modifications in cancer malignancy, we investigated proteins that showed changes in amount and/or modification after the translation process by comparison between western blotting and the microarray expression data to discover the factors that enhanced tumorigenicity. We employed LM1-2-1 cells as the comparison subject to search for specific factors in LM05 cells. The analysis results showed that the amounts of mRNA and protein related with EGFR-MAPK, AKT-mTOR and JAK-STAT pathways were not changed (Fig. 1C, D), although the protein level of NF-κB2 (p52) increased (Fig. 1D). The activation of NF-κB signaling results in nuclear translocation of the NF-κB family of proteins: p105/p50 (NF-κB1), p100/p52 (NF-κB2), RelA (p65), and RelB ^20,21^. Therefore, we investigated the nuclear translocation of the NF-κB family proteins by western blotting to evaluate the activity of NF-κB signaling in LM05 cells. Signaling analysis revealed that NF-κB2 (p52) and RelB increased in the nuclear fraction of LM05 cells compared with the parental cells and LM1-2-1 cells (Fig. 1E). On the other hand, p50 and RelA in the nuclear fraction were unchanged in LM05 cells (Fig. 1E). Since p52 and RelB are classified as part of the non-canonical NF-κB pathway, we examined the amounts of NF-kappa-β-inducing kinase (NIK), a key regulator of the non-canonical NF-κB pathway. NIK is rapidly degraded by ubiquitin-proteasome system after ubiquitinated with TRAF2-TRAF3-cIAP1/2 complex ^22,23^. Therefore, we investigated the amounts of NIK protein in the presence of a proteasome inhibitor, MG132. As a result, NIK was found to be increased at the protein level even though NIK mRNA expression was not significantly changed in LM05 cells (Fig. 1F). These results indicated that the non-canonical NF-κB pathway was constitutively activated in LM05 cells via NIK upregulation at the protein level.

**Figure 1.**
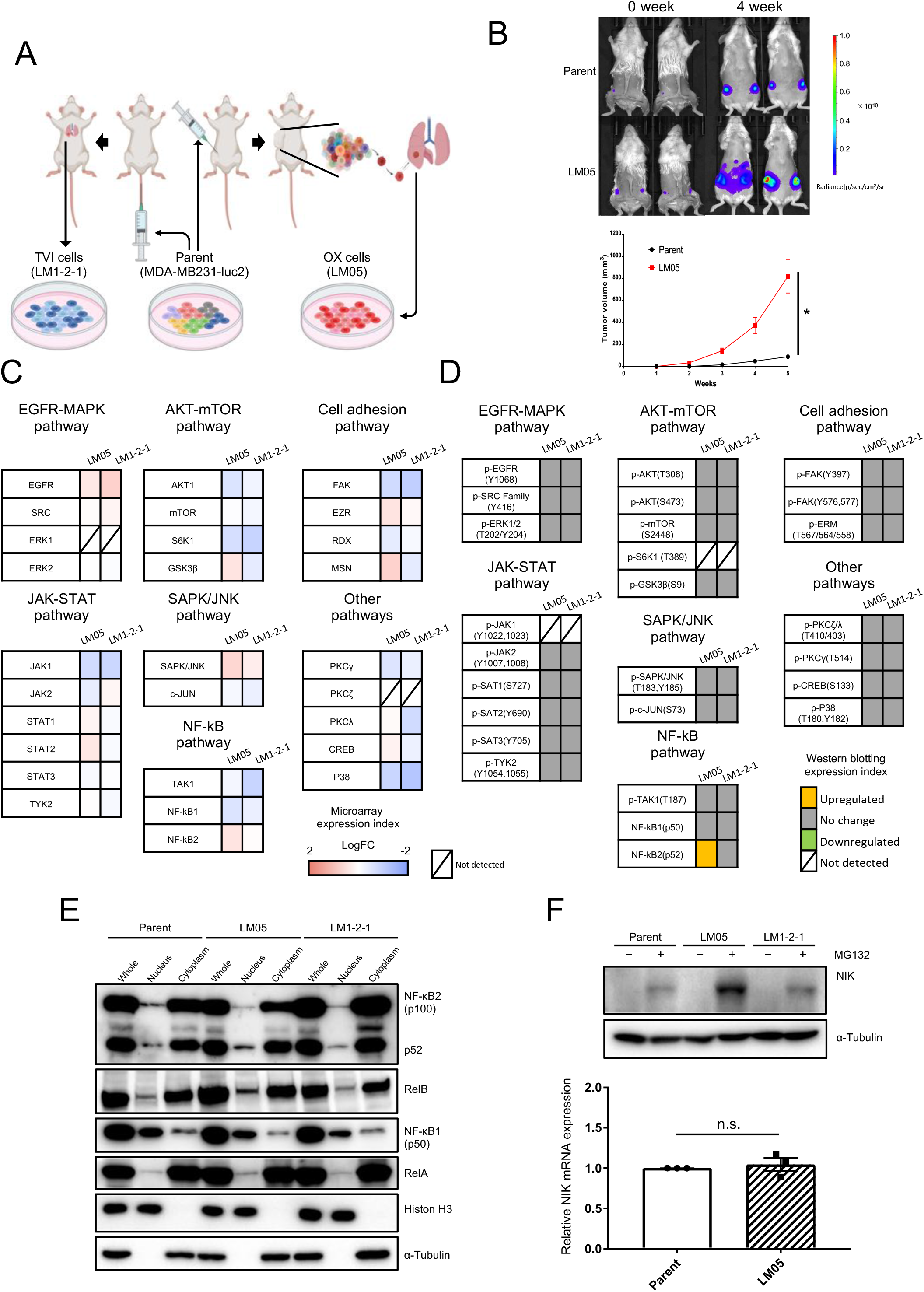
The non-canonical NF-κB pathway was constitutively activated in LM05 cells via upregulation of NIK at the protein level. **A.** Schematic representation of the *in vivo* selection process using an orthotopic xenograft (OX) and tail vein injection (TVI). Luciferase-expressing MDA-MB-231 (parental cells) cells were transplanted into NOD-SCID mice by OX or TVI. Subsequently, lung metastatic cells were collected and established from lung tissue with metastases. These cells were reinjected into NOD-SCID mice using the same xenograft model to concentrate these cells with higher lung metastatic activity. **B.** Representative *in vivo* bioluminescent images (upper) and tumor growth curves (lower, n=4 per group, two-way ANOVA followed by Tukey’s multiple comparison test) of NOD-SCID mice orthotopically injected with parental or LM05 cells at 0 week and 5 weeks. **C.** Integrative signaling analysis of LM05 cells analyzed by microarray expression data. The microarray expression data results are shown as a heatmap of the log_2_-fold change in each cell compared with the parental cells. **D.** Integrative signaling analysis of LM05 cells analyzed by western blotting. The western blotting results had four evaluation criteria (yellow: upregulated, green: downregulated, and gray: unchanged protein production levels compared with parental cells, diagonal line: undetected protein). **E.** Western blotting analysis of NF-κB1 (p50), NF-κB2 (p100/52), RelA and RelB in whole cells and the nuclear and cytoplasmic extracts of the parental, LM05, and LM1-2-1 cells. **F.** Western blotting (upper) and qRT-PCR (lower, n=3, Welch’s t test) results of NIK protein and mRNA in the parental, LM05, and LM1-2-1 cells. For western blotting, all cell lines were either untreated or treated with MG132 (10 μM for 4 hr). All data are representative of three independent experiments and are shown as the mean±SEM. NS, not significant. * P<0.05.

### NIK upregulation was induced by suppression of nascent cIAP1 protein production

NIK is constantly degraded in the normal state by the TRAF2-TRAF3-cIAP1/2 complex in the ubiquitin-proteasome system ^22,23^. We therefore examined the ubiquitination of NIK by immunoprecipitation with a NIK antibody. Ubiquitination of NIK was found to be decreased in LM05 cells compared with parental cells (Fig. 2A). Next, we examined the amounts of cIAP1, which acts as an E3 ligase to ubiquitinate NIK ^22,23^. cIAP1 was found to be decreased at the protein level even though cIAP1 mRNA expression was unchanged compared to the parental and LM05 cells (Fig. 2B). Notably, NIK upregulation in LM05 cells was suppressed by cIAP1 overexpression (Supplementary Fig. S1A). From these results, the mechanism of NIK upregulation in LM05 cells was dependent on cIAP1 reduction. Next, we evaluated the protein stability of cIAP1 by cycloheximide chase analysis. These results demonstrated that the cIAP1 reduction at the protein level was not ascribed to protein degradation in LM05 cells (Fig. 2C). In previous studies, it was established that nascent cIAP1 production was occasionally regulated by translational mechanisms ^24–26^. Hence, we evaluated the production of the nascent cIAP1 protein pulse-labeled with L-homopropargylglycine (HPG) as a methionine analog conjugated to a biotin tag using a click reaction. The results showed that HPG-labeled cIAP1 protein production decreased in LM05 cells compared with parental cells (Fig. 2D). All of these results indicated that the downregulation of cIAP1 translation caused NIK stabilization, which contributed to the constitutive activation of the non-canonical NF-κB pathway.

**Figure 2.**
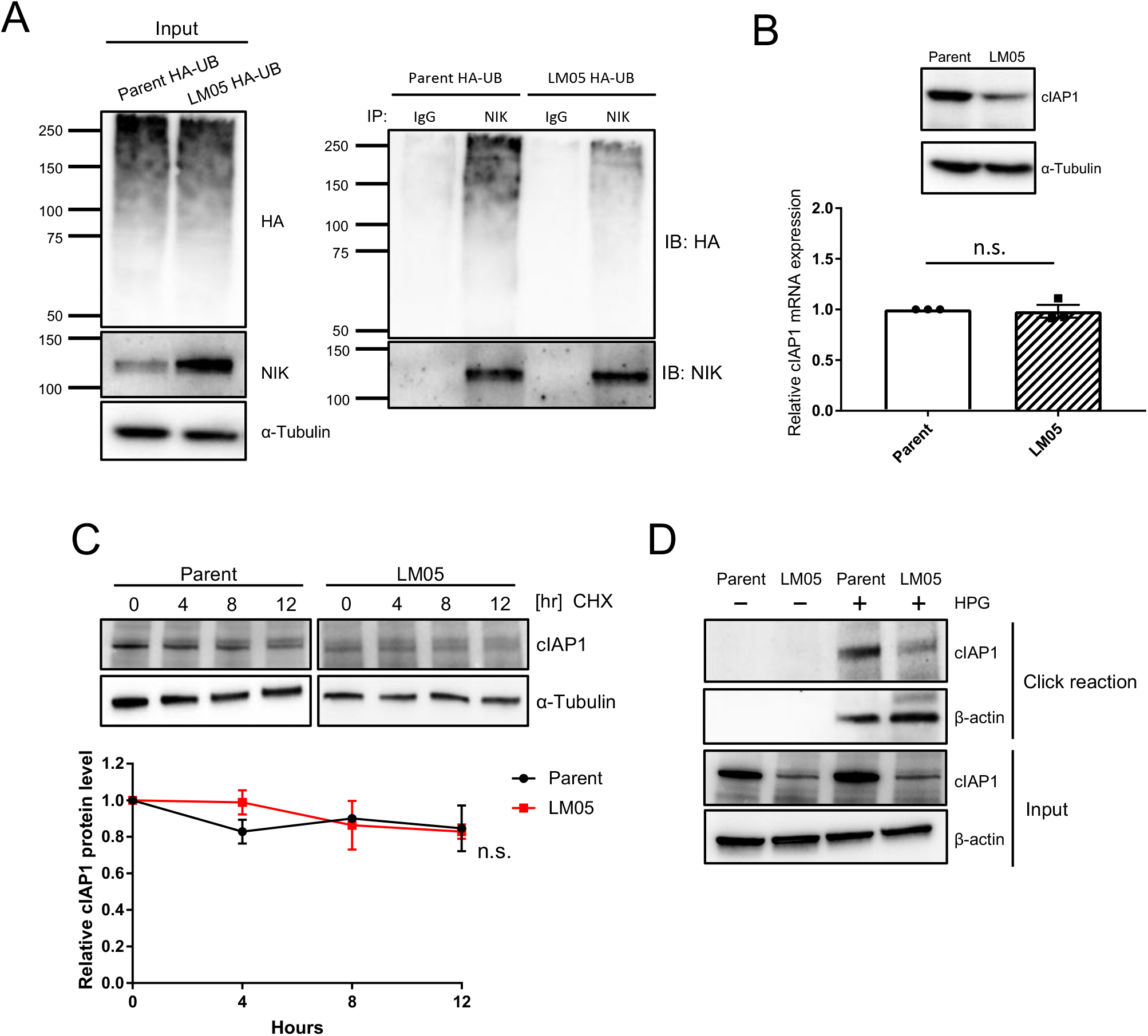
Downregulation of cIAP1 translation caused NIK stabilization, which contributed to constitutive activation of the non-classical NF-κB pathway. **A.** Western blotting analysis of ubiquitinated NIK in HA-tagged ubiquitin (HA-UB)-expressing parental or LM05 cells after treatment with 10 μM MG132 for 4 hr. The protein lysates of HA-UB expressed in parental or LM05 cells were immunoprecipitated with an anti-NIK antibody and then immunoblotted with an anti-HA antibody. **B.** Western blotting (upper) and qRT-PCR (lower, n=3, Welch’s t test) analysis of cIAP1 expression in parental and LM05 cells. **C.** Assessment of cIAP1 protein stability in parental and LM05 cells treated with 10 µg/mL cycloheximide (CHX). Quantification of cIAP1 protein level normalized by α-Tubulin protein level in each time point (n=3, two-way ANOVA followed by Bonferroni’s multiple comparisons test). **D.** Assessment of nascent cIAP1 protein levels in parental and LM05 cells using a click reaction. Parental and LM05 cells were treated with 50 μM L-homopropargyl glycine (HPG) for 24 hr. The nascent proteins labeled with HPG were conjugated to biotin using a click reaction. The biotinylated proteins were purified with streptavidin beads and subjected to western blotting. All data are representative of three independent experiments and are shown as the mean±SEM. NS, not significant.

### NIK knockdown reduced colony formation activity in LM05 cells

To determine the contribution of NIK to cancer malignancy, we characterized the phenotype of LM05 cells after NIK knockdown *in vitro* (Fig. 3A). NIK knockdown cells depressed the non-canonical NF-κB pathway by suppressing the nuclear localization of p52 and RelB (Fig. 3B). We next evaluated cell proliferation and anchorage-independent growth with a soft agar assay. NIK expression did not affect cell growth in planar culture (Fig. 3C), although the colony formation activity was decreased by NIK knockdown (Fig. 3D). We therefore assessed the contribution of NIK knockdown to cancer stemness, since NIK is known to be a supportive factor of cancer stemness from the results of previous studies ^27,28^. Notably, NIK knockdown did not bring about changes in the stemness population rate (Fig. 3E) or sphere formation in the mammosphere culture assay (Fig. 3F). From these results, NIK upregulation in LM05 cells contributed to colony formation but not cancer stemness.

**Figure 3.**
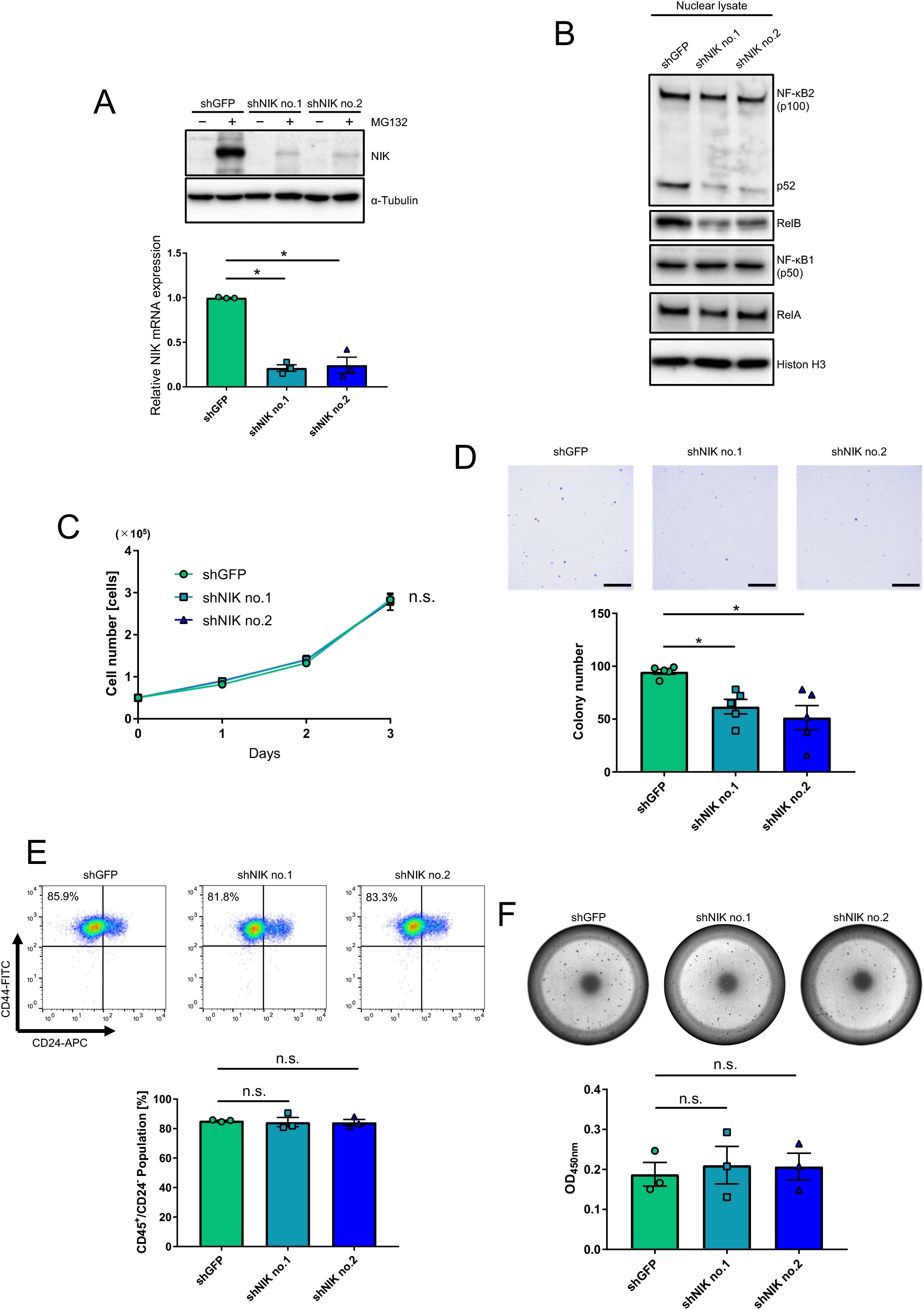
NIK upregulation in LM05 cells contributed to colony formation activity but not cancer stemness. **A.** Validation of NIK knockdown efficiency via Western blot (upper) and qRT-PCR (lower, n=3, one-way ANOVA followed by Tukey’s multiple comparison test) analyses in LM05-shGFP, shNIK no.1 and shNIK no.2 cells. For western blotting, all cell lines were either untreated or treated with MG132 (10 μM for 4 hr). **B.** Western blotting analysis of NF-κB1 (p50), NF-κB2 (p100/52), RelA and RelB in the nuclear extracts of LM05-shGFP, shNIK no.1 and shNIK no.2 cells. **C.** Cell growth curves of LM05-shGFP, shNIK no.1 and shNIK no.2 cells from planar culture (n=3, one-way ANOVA followed by Tukey’s multiple comparison test). **D.** Representative images (upper) and quantification data (lower) (n=5, one-way ANOVA followed by Tukey’s multiple comparison test) of the soft agar colony formation assay in LM05-shGFP, shNIK no.1 and shNIK no.2 cells. The scale bar is 5mm. **E.** Representative images (upper) and quantification data (lower) (n=3, one-way ANOVA followed by Tukey’s multiple comparison test) from the flow cytometry analysis of the cancer stem cell populations in LM05-shGFP, shNIK no.1 and shNIK no.2 cells. **F.** Representative images (upper) and quantification data (lower) (n=3, one-way ANOVA followed by Tukey’s multiple comparison test) of the mammosphere culture assay in LM05-shGFP, shNIK no.1 and shNIK no.2 cells. All data are representative of three independent experiments and are shown as the mean±SEM. NS, not significant. * P<0.05.

### Upregulation of NIK contributed to tumorigenicity in LM05 cells

To investigate whether NIK upregulation contributes to the inherent high tumorigenicity and potential of LM05 cells to metastasize to the lungs, NIK knockdown cells were orthotopically injected into the mammary fat pats of NOD-SCID mice. NIK knockdown in LM05 cells significantly decreased the primary tumor weights and volumes (Fig. 4A and B). In addition, the reduction in tumorigenicity caused by NIK knockdown was partially rescued by recovering the ectopic expression of NIK (Supplementary Fig. S2A, B and C). In addition, we examined the effects of NIK knockdown on metastatic potential by performing a metastasis assay in an orthotopic xenograft model. To remove the effects from tumorigenicity reduction created by NIK knockdown, we uniformly resected the primary tumors after they reached a size of 300 mm^3^. Under these conditions, NIK knockdown did not affect the potential of LM05 cells to form lung metastases (Fig. 4C). Furthermore, IHC analysis of lung metastases using an anti-CAM5.2 antibody, a human-specific cytokeratin, indicated that there were no differences in the metastatic node areas between the control and NIK knockdown groups (Fig. 4D). These results indicated that NIK upregulation facilitated the inherent tumorigenicity but not lung metastatic potential of LM05 cells.

**Figure 4.**
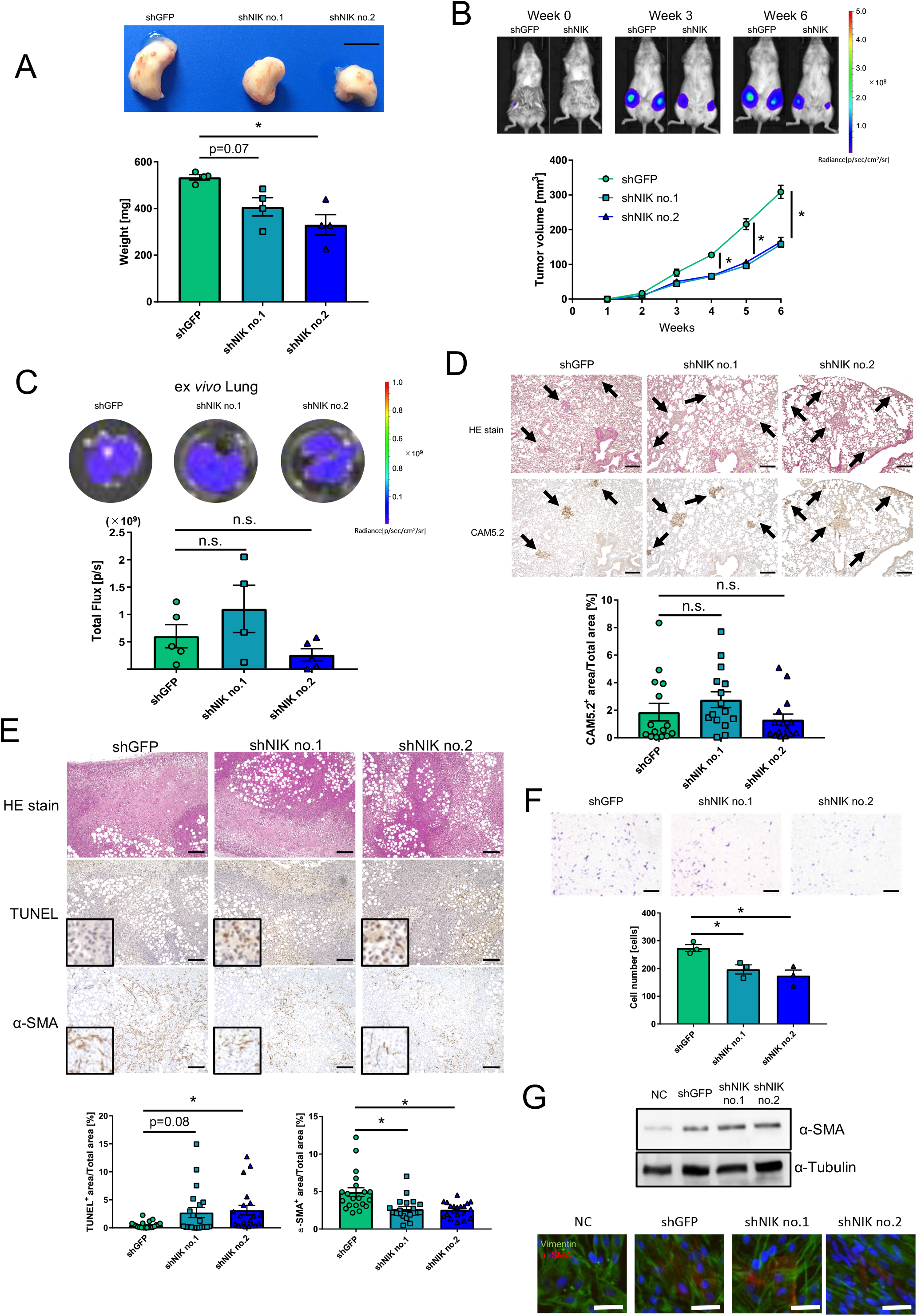
NIK upregulation facilitated inherent tumorigenicity but not the lung metastatic potential of LM05 cells. **A.** Representative images of primary tumors (upper) and quantification data of the primary tumor weights (lower) (n=4, one-way ANOVA followed by Tukey’s multiple comparison test) in LM05-shGFP, shNIK no.1 and shNIK no.2 cells. The scale bar is 1cm. **B.** Representative *in vivo* bioluminescent images of LM05-shGFP and shNIK no.2 cells (upper). Tumor growth curves (lower) (n=6 per group, two-way ANOVA followed by Tukey’s multiple comparison test) of NOD-SCID mice orthotopically injected with LM05-shGFP, shNIK no.1 and shNIK no.2 cells. **C.** Representative *in vivo* bioluminescent images (upper) and quantification data of the lung metastasis tissue (lower) (one-way ANOVA followed by Tukey’s multiple comparison test) derived from NOD-SCID mice orthotopically injected with LM05-shGFP (n=5), shNIK no.1 (n=4) and shNIK no.2 cells (n=5). **D.** Representative HE and IHC staining images (upper) and quantification data (lower) (one-way ANOVA followed by Tukey’s multiple comparison test) of NIK protein production in lung metastasis tissue derived from NOD-SCID mice orthotopically injected with LM05-shGFP, shNIK no.1 and shNIK no.2 cells. The average CAM5.2-positive percentages were calculated from 5 fields of view from n=4 individual lung metastasis slides. The scale bar is 500 μm. **E.** Representative HE and IHC staining images (upper) and quantification data (lower) (one-way ANOVA followed by Tukey’s multiple comparison test) from the TUNEL assay and α-SMA protein production in primary tumor tissues derived from NOD-SCID mice orthotopically injected with LM05-shGFP, shNIK no.1 and shNIK no.2 cells. The average number of TUNEL- and α-SMA-positive cells were calculated from 5 fields of view from n=4 individual primary tumor slides. The scale bar is 500 μm. **F.** Representative images (upper) and quantification data (lower) (n=5, one-way ANOVA followed by Tukey’s multiple comparison test) of the Boyden chamber assay with TIG-3 cells that were co-cultured with LM05-shGFP, shNIK no.1, and shNIK no.2 cells. The scale bar is 500 μm. **G.** Western blotting (upper) and immunofluorescence staining (lower) of α-SMA expression in TIG-3 cells that were co-cultured with LM05-shGFP, shNIK no.1 and shNIK no.2 cells or noncells (NC). The scale bar is 50 μm. All data are shown as the mean±SEM. NS, not significant. * P<0.05.

Next, we performed immunohistochemical staining of the primary tumor to investigate the role of NIK in the primary tumor. The number of TUNEL-positive cells (apoptosis marker) increased in the primary tumor after NIK knockdown (Fig. 4E). In addition, the α-SMA-positive region (a cancer-associated fibroblast (CAF) marker) decreased in the primary tumors after NIK knockdown (Fig. 4E). Therefore, we utilized a co-culture system in a Boyden chamber to investigate the effects of NIK on the human fibroblast cell line TIG-3. From the experimental results, it was found that NIK knockdown suppressed the attraction of TIG-3 cells to the cancer cells in the co-culture system (Fig. 4F). On the other hand, α-SMA protein in TIG-3 cells did not change when co-cultured with control and NIK knockdown cells (Fig. 4G), indicating that NIK is involved in attraction of fibroblasts but not induce their activation to CAFs. These results suggest that the suppression of tumorigenesis caused by NIK knockdown is due to increased apoptosis of the tumor cells and decreased induction of CAFs.

### NIK knockdown suppressed cancer-inducing inflammatory signaling

To determine the NIK-related pathway and genes that contribute to the phenotypes altered by NIK knockdown (Fig. 4A, B and E), we performed transcriptome analysis by RNA-seq. We extracted the differentially expressed genes (DEGs) between LM05-shGFP cells and LM05-shNIK cells using the following criteria: FDR <0.05 (Benjamini-Hochberg method) and log fold change (Log_2_(FC)) ±1 (Fig. 5A). To determine the NIK-related pathways, the common DEGs in the NIK knockdown cells were analyzed by gene set enrichment analysis (GSEA) and ingenuity pathway analysis (IPA). GSEA using the MSigDB Hallmark gene set collection revealed that the genes with decreased expression in the NIK knockdown cells were significantly enriched for pathways related to inflammation, interferon response, and TNFα signaling via NF-κB (Fig. 5B, Supplementary Table S2). Similarly, pathways related to the interferon response and TNF secretion were suppressed in NIK knockdown cells using Gene Ontology gene sets (Supplementary Table S3). Furthermore, upstream regulator analysis by IPA contained components of NF-κB signaling as an upstream regulator of the downregulated pathway (Supplementary Fig. S3A). These transcriptome analyses showed that the inflammatory-related genes regulated by the NF-κB pathway were suppressed in LM05 NIK knockdown cells. Indeed, NIK knockdown decreased the expression of *IL6* and *CXCL1*, which are important for the migration of fibroblasts (Fig. 5C) ^29,30^. Furthermore, *BIRC3* expression, which contributes to apoptosis resistance, was markedly reduced by NIK knockdown (Fig. 5C) ^31,32^. Based on these results, the reduction in *BIRC3* expression and fibroblast-attracted cytokines and chemokines caused by NIK knockdown inherently result in decreased tumorigenicity.

**Figure 5.**
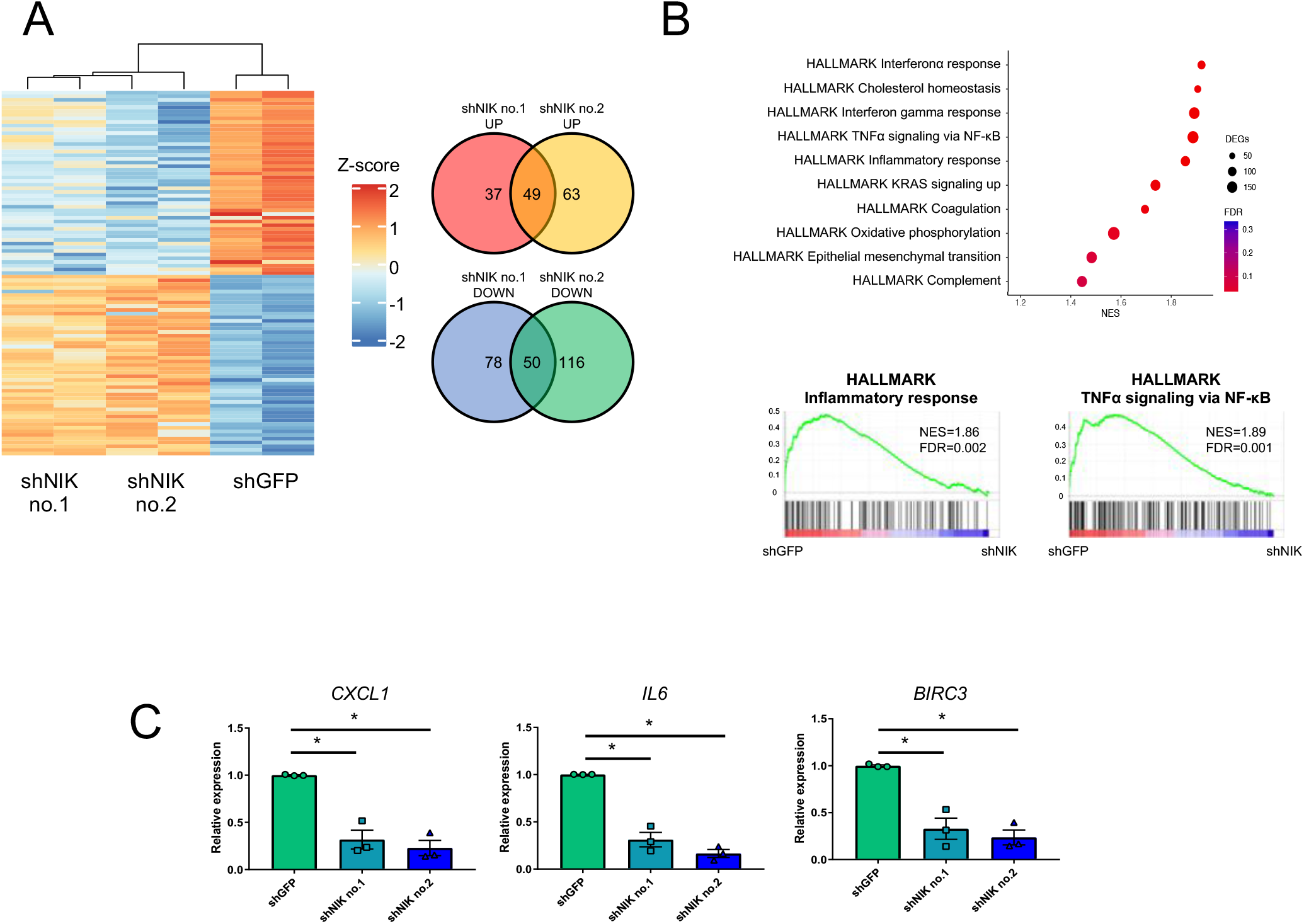
NIK regulated cancer-inducing inflammatory signaling in LM05 cells. **A.** Heatmap and Venn diagram of the DEGs from the LM05-shGFP, LM05-shNIK no.1 and LM05-shNIK no.2 RNA-seq data, each performed in duplicate. Hierarchical analysis of the heatmap was performed by the complete linkage method. Venn diagram of the signature genes in LM05-shNIK cells. **B.** GSEA enrichment plot of the hallmark gene sets for the differentially expressed genes in LM05-shNIK cells compared with LM05-shGFP cells. **C.** Validation of NIK signature gene expression related to tumor inflammation using qRT-PCR (lower) (n=3, one-way ANOVA followed by Tukey’s multiple comparison test) in LM05-shGFP, shNIK no.1 and shNIK no.2 cells. All data are shown as the mean±SEM. NS, not significant. * P<0.05.

### NIK protein production increased in malignant breast cancer tissue

We then investigated the relationship between NIK production and breast cancer malignancy. We characterized NIK production by immunohistochemistry in a tissue microarray from breast cancer patients (Fig 6A). IHC analysis showed that NIK production was significantly increased in stage I-III patients compared with normal breast tissues (Supplementary Table S4, Fig. 6B). In addition, NIK protein production was not correlated with the expression scores of breast cancer marker genes such as ER, PR, and HER2 (Supplementary Fig. S4A, B and C). From these results, NIK has the potential to be used as a diagnostic marker across other breast cancer subtypes.

**Figure 6.**
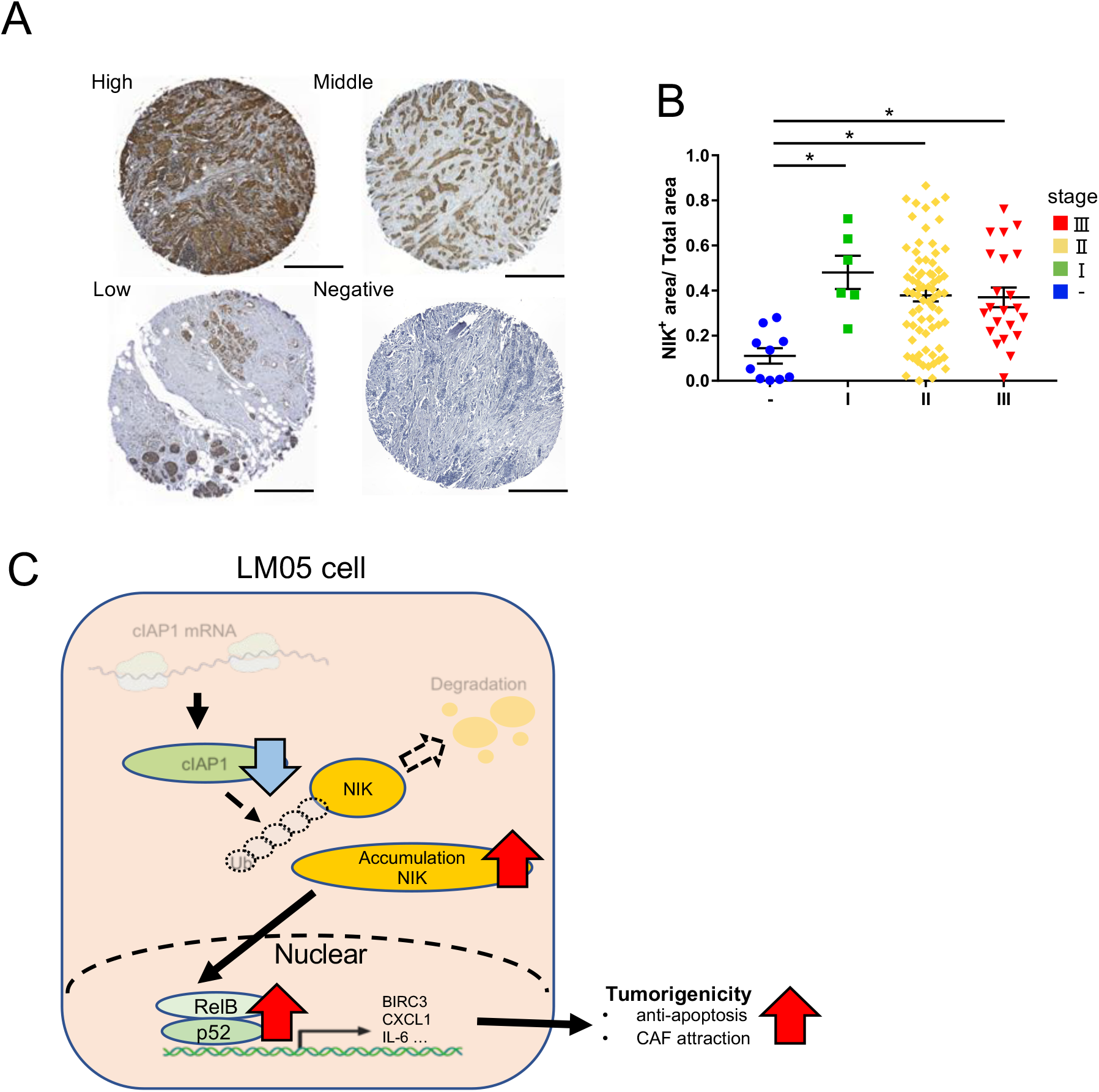
NIK protein production increased in malignant breast cancer tissue. **A.** Representative IHC staining images of NIK protein production in normal breast tissue and breast tumors. The scale bar is 500 μm. **B.** Quantification data of the NIK IHC staining images in normal breast tissue and breast tumors ((n=10 normal, n=6 stage I, n=72 stage II, and n=22 stage III); one-way ANOVA followed by Tukey’s multiple comparison test). **C.** The graphical summary indicates that the abnormal accumulation of NIK due to reduced translation of cIAP1 enhances tumorigenesis by promoting an inflammatory cancer microenvironment. All data are shown as the mean±SEM. NS, not significant. * P<0.05.

## Discussion

Constitutive activation of the non-canonical NF-κB pathway has been reported in breast cancer cell lines and multiple myeloma cell lines ^33–35^. In breast cancer, basal-like subtype cell lines have shown increased NIK mRNA expression by epigenetic dysregulation of NIK gene, which induces constitutive activation of the non-canonical pathway ^35,36^. On the other hand, certain multiple myeloma cell lines have genetic defects in TRAF2/3 and cIAP1/2 ^33,34^, which are repressors of the non-canonical pathway. However, constitutive activation of the non-canonical pathway due to reduced translation of cIAP1, as revealed in this study, is a novel activation mechanism of the non-canonical pathway in breast cancer cell lines (Fig. 6C). In previously, the activation of the non-canonical NF-κB pathway by NIK upregulation contributes to the enhancement in cell proliferation and maintenance of cancer stemness properties via the NOTCH pathway, especially in basal-like breast cancer cell lines ^27,35^. Although claudin-low cell lines exhibit high levels of NF-κB activation with increased NIK mRNA expression ^35,36^, the function of NIK in tumorigenesis and metastasis has not been investigated. Our results suggested that NIK upregulation contributed to tumorigenicity by attracting CAFs and inhibiting apoptosis through regulation of the expression of genes related to inflammatory responses in LM05 cells, which is classified as the claudin-low subtype (Fig. 6C). Therefore, there are novel function and aberrant accumulation mechanism of NIK in breast cancer malignancy that differs from the conventional researches.

From our results, a reason for the reduction of tumorigenicity by NIK knockdown is the expansion of the apoptotic area in the primary tumor. In addition, our results suggest that the decrease in *BIRC3* expression, which has antiapoptotic properties, and CAFs may contribute to the increase in the apoptotic area. CAFs have been reported to contribute to the enhancement of tumor malignancy by promoting the antiapoptotic potential and chemoresistance of cancer cells ^37,38^. It is possible that a reduction in the number of CAFs in tumors by NIK knockdown contributes to the decline in the antiapoptotic potency of cancer cells. CXCL12/SDF-1 secreted by CAFs contributes to tumor growth and anti-apoptosis in cancer cells via CXCR4 ^39,40^. Indeed, the expression of *CXCR4* was reduced in NIK knockdown cells; thus, there was a deficiency in the positive feedback through this paracrine mechanism. Therefore, NIK plays an important role in the interaction with stromal cells in the cancer microenvironment.

NIK is crucial for the maintenance of various tissue functions, including the immune system, bone formation, the kidneys, the liver, glucose homeostasis and hematopoiesis ^41^. Therefore, abnormal NIK activation has been implicated in a variety of autoimmune diseases, such as systemic lupus erythematosus, acute kidney injury and cancer ^41^. Recently, exploratory studies of NIK inhibitors have expanded ^41^. NIK inhibitors have been developed by modifying inhibitors of similar kinase family members and conventional NIK inhibitors based on structure-activity relationships and docking simulations using NIK crystallographic data ^42,43^. These studies have identified that NIK inhibitors have therapeutic effects in certain mouse models of inflammatory hepatic diseases and systemic lupus erythematosus ^44,45^. In addition, mangiferin, which is a natural compound with NIK inhibitory activity, has been reported to inhibit tumorigenesis and metastatic potential in melanoma cell lines ^46^. However, the efficacy of these NIK inhibitors against other types of cancer is unclear; thus, verification of the activity of these NIK inhibitors is needed to examine this possibility. In contrast, inactivating NIK mutations have been shown to cause immunodeficiency, such as decreased numbers of mature B cells and T cells their functional impairment ^47^. NIK plays an important role in antitumor immunity by regulating metabolism in cytotoxic CD8^+^ T cells ^48^. Therefore, it is necessary to develop NIK inhibitors through further comprehension of the reciprocal NIK functions to avoid interfering with the homeostatic role that NIK plays in the body. Our experimental results indicated that NIK knockdown critically affected tumorigenicity rather than the potential to form lung metastases in LM05 cells. From this result, we hypothesized that NIK may play an important role in tumorigenesis mechanisms in the mammary gland. To address this hypothesis, we evaluated the proliferative potential of parental and LM05 cells in lung tissue using tail vein injection. The results showed that there was no significant difference in the proliferative potential of the lung tissues between the parental and LM05 cells (Supplementary Fig. S5A). In addition, LM05 cells were repeatedly xenografted into the mammary gland and grown at this site. Thus, the enhanced tumorigenicity of LM05 is probably mammary gland-specific. Indeed, TWEAK and RANKL, ligands of the non-canonical NF-κB pathway, are expressed and have important functions in mammary gland development and tumorigenesis ^49–52^. In short, LM05 cells may be more susceptible to the activation of the non-canonical pathway by these ligands in breast tissue because of the accumulation of NIK proteins. Meanwhile, we were not able to identify the genes that contribute to the enhanced potential of LM05 cells to metastasize to the lungs. However, we identified a group of genes that were upregulated in LM05 cells compared to the parental strain but not downregulated after NIK knockdown, including known lung metastasis regulatory genes such as *IL13RA2*, *TNS1*, and *EMP1* ^18,53,54^ (Supplementary Fig. S6A). Further studies of those genes may reveal the NIK-independent metastasis enhancement mechanism in LM05 cells.

In conclusion, our results demonstrated a novel role and mechanism of NIK after its upregulation during breast cancer progression. In addition, the amount of NIK protein was increased in tumor tissues compared to normal tissues; thus, NIK has the potential to be a diagnostic marker of breast cancer. Elucidating the functions and regulatory mechanisms of NIK can lead to a deeper understanding of NIK as a potential biomarker or therapeutic target in breast cancer.

## Acknowledgments

We thank Dr. Bunsyo Shiotani (National Cancer Center Research Institute) for providing the TIG-3 cells. We are also grateful to our laboratory member for worthwhile discussion and technical advice. Y.H. was supported by doctoral scholarships of the Futaba foundation. Schematics in figures were made using an academic license of BioRender.com.

## Authors’ contributions

J.N. and K.S. conceived and designed the study. Y.H. and J.N. performed the analyses and the experiments. M.Y., M.M., S.W., S.H., J.I. and Y.Y. interpreted the data. Y.H. wrote the original paper. J.N., Y.Y. and K.S. revised the manuscripts. All authors reviewed and edited the manuscript.

---

**Supplementary Figure 1.**
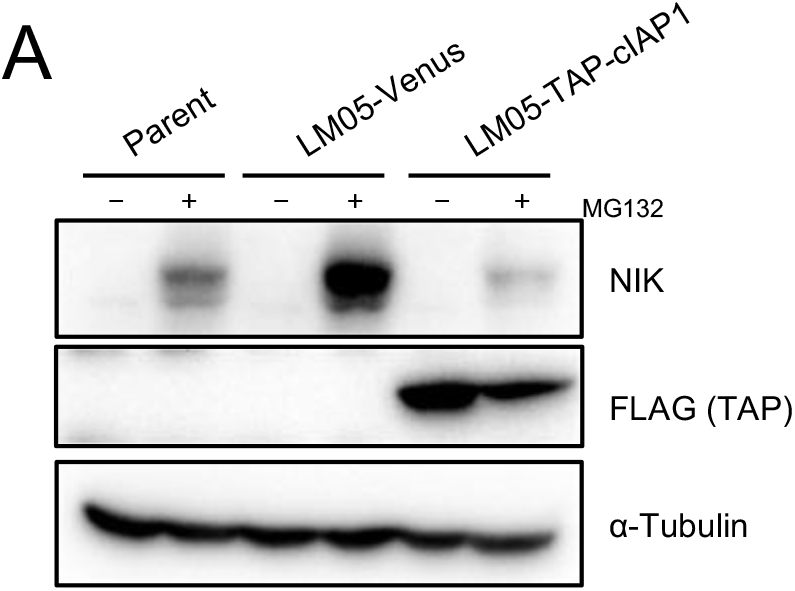
cIAP1 ectopic expression reduced NIK protein production in LM05 cells. Western blotting analysis of NIK protein production in parental, LM05-Venus and LM05-TAP-cIAP1 (murine) cells. For western blotting, all cell lines were either untreated or treated with MG132 (10 μM for 4 hr). All data are representative of three independent experiments.

**Supplementary Figure 2.**
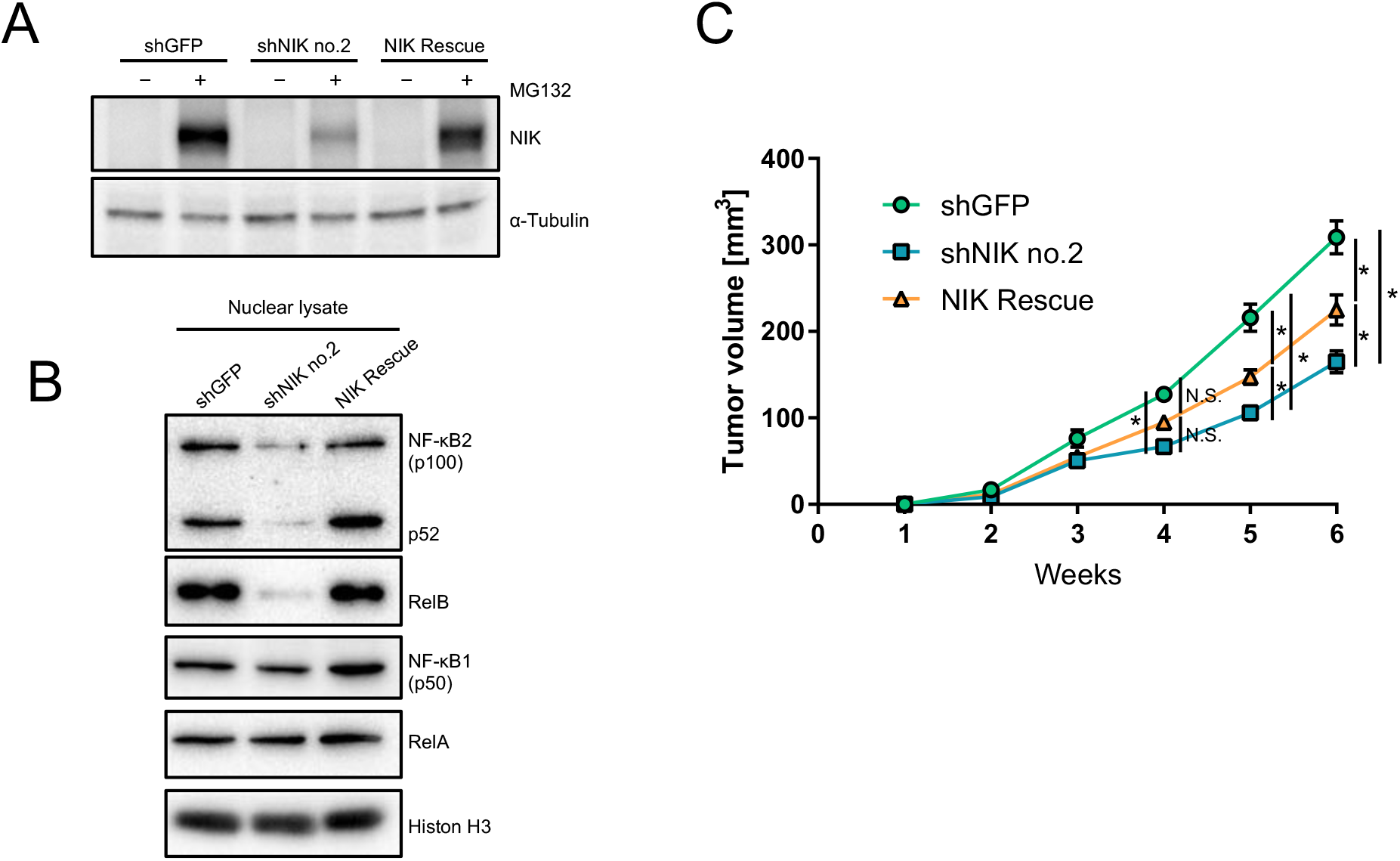
NIK ectopic expression partially rescued the reduction in tumorigenicity induced by NIK knockdown. **A.** Validation of the NIK rescue efficiency via Western blot analysis in LM05-shGFP, shNIK no.2 and NIK rescue cells. For western blotting, all cell lines were either untreated or treated with MG132 (10 μM for 4 hr). **B.** Western blotting analysis of NF-κB1 (p50), NF-κB2 (p100/52), RelA and RelB in the nuclear extracts of LM05-shGFP, shNIK no.2 and NIK rescue cells. **C.** Tumor growth curves (two-way ANOVA followed by Tukey’s multiple comparison test) of NOD-SCID mice orthotopically injected with LM05-shGFP (n=6), shNIK no.2 (n=6) and NIK (n=4) rescue cells. All data are representative of three independent experiments and are shown as the mean±SEM. NS, not significant. * P<0.05.

**Supplementary Figure 3.**
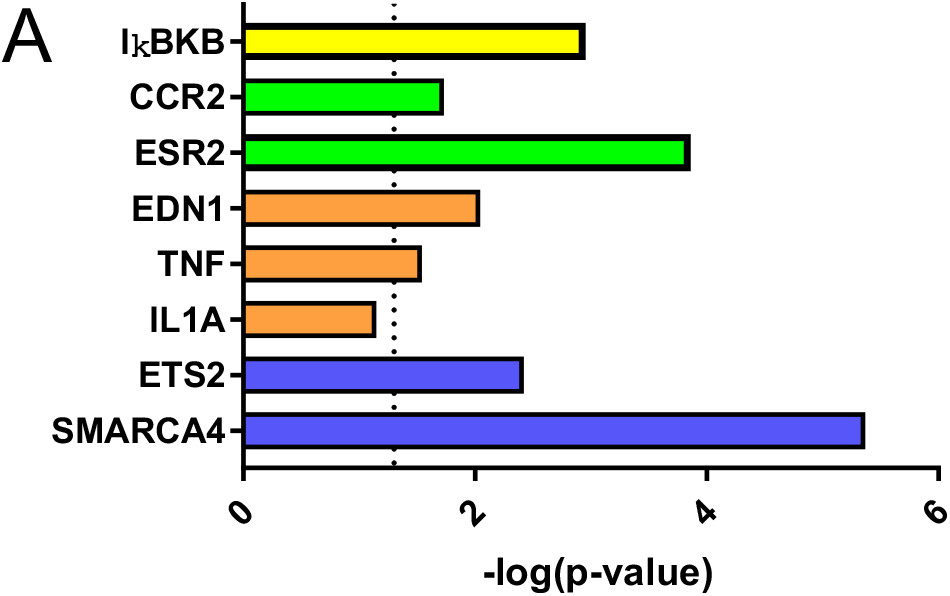
The upstream regulators of the downregulated common DEGs in NIK knockdown cells contained some genes associated with NF-κB signaling. **A.** Ingenuity pathway analysis suggested statistically significant upstream regulators of the downregulated common DEGs in NIK knockdown cells (selected activation z-score < −2). The bar color indicates the molecule type of the upstream regulator (orange: cytokine, blue: transcription regulator, yellow: kinase, green: receptor). The dotted line shows a p-value = 0.05.

**Supplementary Figure 4.**
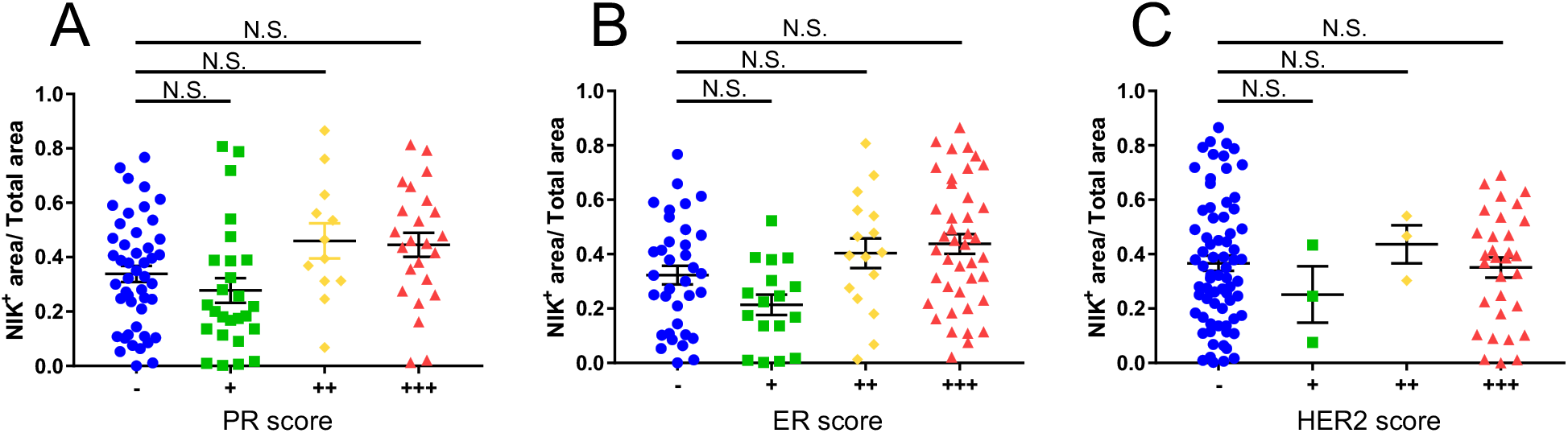
NIK expression was not correlated with PR, ER, or HER2 scores. **A.** Quantification data of the NIK IHC staining images of normal breast tissue and breast tumors for each PR score (n=47 - (negative), n=26 +, n=12 ++, and n=24 +++; one-way ANOVA followed by Tukey’s multiple comparison test). **B.** Quantification data of the NIK IHC staining images of normal breast tissue and breast tumors for each ER score (n=35 - (negative), n=17 +, n=26 ++, and n=41 +++; one-way ANOVA followed by Tukey’s multiple comparison test). **C.** Quantification data of the NIK IHC staining images of normal breast tissue and breast tumor for each HER2 score (n=72 - (negative), n=3 +, n=3 ++, and n=31 +++; one-way ANOVA followed by Tukey’s multiple comparison test).

**Supplementary Figure 5.**
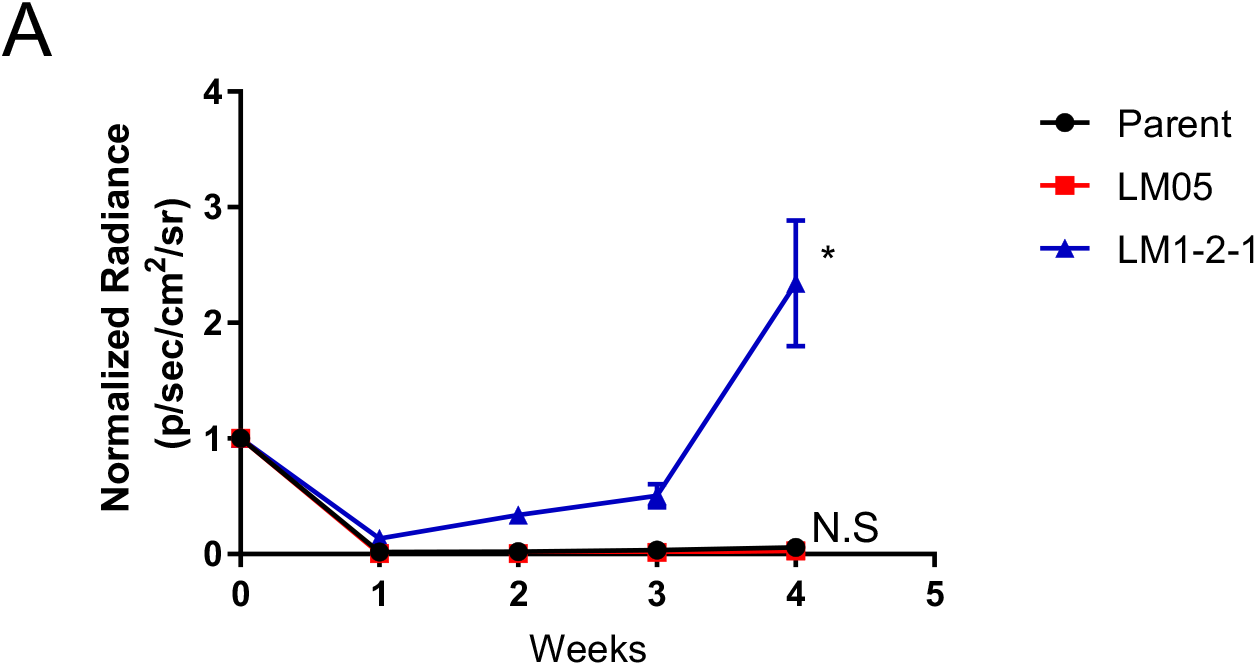
The TVI model showed that the lung metastatic potential of LM05 cells was not enhanced compared with parental cells. **A.** Representative *in vivo* bioluminescent images (left) and quantification data of lung metastases (right) (one-way ANOVA followed by Tukey’s multiple comparison test) derived from NOD-SCID mouse tail vein injection with parental, LM05, or LM1-2-1 cells (n=5).

**Supplementary Figure 6.**
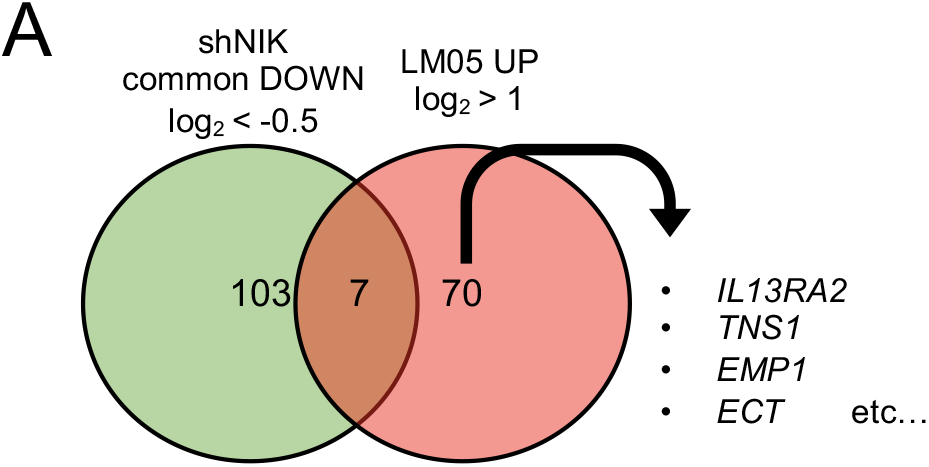
Some known lung metastasis-promoting genes are highly expressed in the LM05 cell line independent of NIK knockdown. **A.** Venn diagram of the upregulated genes in LM05 cells based on previous microarray expression data and the downregulated genes found in the NIK KD cell line based on the RNA-seq data from this study. Expression of the known metastasis-promoting genes *IL13RA2*, *TNS1*, and *EMP1* is independent of NIK knockdown.

