## Supplementray Method for "Aberrant accumulation of NIK promotes tumorigenicity by dysregulating post-translational modifications in breast cancer"

**Supplemental method**

*Expression and knockdown vector constructs*

NIK, cIAP1 and HA tag-fused ubiquitin (HA-UB) expression vectors were previously produced ^1,2^. The coding region of HA-UB was transferred to the pMXs vector using Ligation High (Toyobo Co., Ltd., Osaka, Japan). shNIK knockdown vectors were previously produced ^3^. All cDNA and shRNA sequences were confirmed by sequencing. Cloning primers and target sequences of shRNA are described in Supplementary Table S5.

*Retroviral packaging and infection*

The production of the retrovirus using Plat-E cells was previously described ^4^. Parent-shGFP, Parent-HA-UB, LM05-shNIK, LM05-shGFP, LM05-HA-UB, LM05-Venus, and LM05-TAP-cIAP1 (murine) cells were established by retroviral infection of the pMXs-target gene cDNA-IRES-Puro^R^ overexpression vectors or pSuper-target shRNA-Psv40-Puro^R^ knockdown vectors and selected with 3 µg/ml puromycin (Fujifilm Wako Pure Chemical Corporation). LM05-NIK rescue cells were established by retroviral infection of pMXs-NIK-Psv40-Neo^R^ overexpression vectors with LM05-shNIK no.2 cells and selected with 1 µg/ml G418 (Fujifilm Wako Pure Chemical Corporation).

*RNA extraction and quantitative reverse transcription polymerase chain reaction (qRT-PCR)*

Total RNA was extracted by ISOGEN (Nippon Gene Co., Ltd., Tokyo, Japan) and reverse-transcribed into cDNA by the SuperScript™ First-Strand Synthesis System for RT-PCR (Thermo Fisher Scientific). Quantitative PCR was performed with Thunderbird SYBR qPCR mix (TOYOBO CO., LTD.) and the StepOnePlus real-time PCR system (Applied Biosystems, Foster, CA, USA). Quantification of the relative mRNA expression levels was performed by normalization to the level of β-actin RNA. The oligonucleotide sequences of the qRT-PCR primers are listed in Supplementary Table S6.

*Soft agar assay*

RPMI culture medium containing 0.3% agarose with LM05-shGFP or LM05-shNIK cells (4.0×10^4^ cells/well) over a bottom layer of 0.6% agarose in RPMI culture medium were plated in each well of a 6-well plate and cultured for 3 weeks. Colonies were fixed with 4% paraformaldehyde-PBS (Fujifilm Wako Pure Chemical Corporation) for 1 hour and stained with 0.005% crystal violet solution (Fujifilm Wako Pure Chemical Corporation) for 30 minutes. After removing the overdyed region with Milli-Q water, the colony images were acquired with a digital camera (Nikon Corporation, Tokyo, Japan), and colony numbers were calculated with ImageJ software (National Institutes of Health).

*Flow cytometry analysis*

Dissociated LM05-shGFP or LM05-shNIK cells were stained with anti-CD24-APC (311118, BioLegend, CA, USA) and anti-CD44-FITC (338808, BioLegend) antibodies at 4°C for 30 minutes. After washing with wash buffer (D-PBS(-) containing 0.5% FBS and 0.5 mM EDTA) three times, the stained cells were passed through a 35 μm cell strainer (Corning, NY, USA) to be dissociated into single cells. Flow cytometry analysis was carried out with an S3e Cell Sorter (Bio-Rad Laboratories, Inc., CA, USA), and data were analyzed with FlowJo software (Becton, Dickinson and Company, NJ, USA).

*Mammosphere assay*

MammoCult medium (MammoCult Basal Medium, MammoCult Proliferation Supplements, containing 4 μg/mL heparin, 0.48 μg/mL hydrocortisone, 0.5% methyl cellulose) (VERITAS Corporation., Tokyo, Japan) was included with LM05-shGFP or LM05-shNIK cells (1.5×10^4^ cells/well) plated on ultralow attachment 24-well plates and cultured for 7 days. To evaluate proliferation under mammosphere culture conditions, Cell Counting Kit-8 reagent (DOJINDO Laboratories, Kumamoto, Japan) was added to the MammoCult medium and incubated for 4 hr at 37°C. After that, the absorbance was measured at 450 nm using a multimode plate reader (Molecular Devices, Tokyo, Japan).

*Migration assay*

LM05-shGFP or LM05-shNIK cells (1.5×10^4^ cells/well) were plated on lower chamber of 24-well plates and incubated for overnight. The culture medium was replaced with Advanced RPMI1640 medium (Thermo Fisher Scientific) supplemented with 1%(V/V) antibiotic-antimycotic (Thermo Fisher Scientific). After incubation for 48hr, the upper chamber of 8μm pore size transwell inserts (Coring), which placed with TIG-3 cells (1×10^4^ cells/well) in serum free medium, placed in the lower chamber. After incubation for 12hr, migrated cells at the lower surface of the membrane were fixed by 4% paraformaldehyde–PBS (Fujifilm Wako Pure Chemical Corporation) and stained by 0.005% crystal violet solution. Then margin liquid and cells on upper surface of the membrane were removed with cotton swab. Images were acquired with a BZ-X700 microscope (Keyence Corporation, Osaka, Japan) and analyzed using ImageJ software (National Institutes of Health).

*Co-culture*

TIG-3 cells (5×10^3^ cells/well) were plated on lower chamber of 6-well plates and incubated for overnight. After incubation, the upper chamber of 0.8 μm pore size transwell inserts (Coring), which placed with LM05-shGFP or LM05-shNIK cells (1×10^4^ cells/well) cells placed in the lower chamber. After incubation for 7days, TIG-3 cells at the lower chamber were lysed in 1× SDS sample buffer and boiling at 95 °C for 5 min, followed by western blotting. In case of immunofluorescence, TIG-3 cells were fixed with 4% paraformaldehyde–PBS for 15min and permeabilized with 0.1% Triton X-100-PBS for 15min. After blocking with Blocking One regent for 1hr, TIG-3 cells were incubated with primary antibodies against α-SMA (#19245, Cell Signaling Technology), Vimentin (V2258, Sigma-Aldrich Co.) at 4°C overnight. Then, TIG-3 cells were incubated with Hoechst 33342 (H3570, Invitrogen, MA, USA) and the secondary antibody conjugated Alexa Fluor 488 and 594 (A11001, A21207, Invitrogen) for 1 hour at room temperature. Images were acquired with a BZ-X700 microscope (Keyence Corporation).

*Click reaction*

Parental and LM05 cells were washed once with D-PBS(-) and incubated for 4 hours in methionine-free RPMI 1640 (Sigma-Aldrich Co.) supplemented with 10% heat-inactivated FBS, GlutaMAX (Thermo Fisher Scientific), and 55 μg/mL L-cystine (Sigma-Aldrich Co.). Then, the media was replaced with media supplemented with 50 μmol/L L-homopropargyl glycine (HPG) (Cayman Chemical, MI, USA) for 24 hours. After the treated cells were dissolved in lysis buffer (50 mM HEPES-KOH (pH 7.3), 150 mM NaCl, 0.1% NP-40, protease inhibitor cocktail), nascent HPG proteins and biotin-PEG3-azide (Tokyo Chemical Industry Co. Tokyo, Japan) were chemically reacted overnight using the Click-iT Protein Reaction Buffer Kit (Thermo Fisher Scientific) at 4°C according to the manufacturer's protocol. After the click reaction, the biotinylated proteins were resuspended in TNE buffer, and Dynabeads™ M-280 Streptavidin (Thermo Fisher Scientific) was added to the resuspended protein lysate. Then, the solution was incubated overnight at 4°C using a rotary shaker. Streptavidin–biotin conjugates were washed five times with TNE buffer. After washing, the conjugates were resuspended in 1× SDS sample buffer and eluted by boiling at 95 °C for 5 min, followed by western blotting.
